# Universal cancer tasks, evolutionary tradeoffs, and the functions of driver mutations

**DOI:** 10.1101/382291

**Authors:** Jean Hausser, Pablo Szekely, Noam Bar, Anat Zimmer, Hila Sheftel, Carlos Caldas, Uri Alon

## Abstract

Recent advances have led to an appreciation of the vast molecular diversity of cancer. Detailed data has enabled powerful methods to sort tumors into groups with benefits for prognosis and treatment. We are still missing, however, a general theoretical framework to understand the diversity of tumor gene-expression and mutations. To address this, we present a framework based on multi-task evolution theory, using the fact that tumors evolve in the body, and that tumors are faced with multiple tasks that contribute to their fitness. In accordance with the theory, we find that tradeoff between tasks constrains tumor gene-expression to a continuum bounded by a polyhedron. The vertices of the polyhedron are gene-expression profiles each specializing in one task, allowing the tasks to be identified. We find five universal cancer tasks across tissue-types: cell-division, biomass & energy, lipogenesis, immune-interaction and invasion & tissue remodeling. Tumors whose gene-expression lies close to a vertex are task specialists. We find evidence that such specialists are more sensitive to drugs that interfere with this task. We find that driver mutations, but not passenger mutations, tune gene-expression towards specialization in specific tasks. This approach can integrate additional types of molecular data into a theoretically-based framework for understanding tumor diversity.

## Introduction

Tumors show diversity in genetic alterations, gene expression and drug sensitivities, presenting a major medical challenge. Tumors continually evolve *(Nowell, 1976; Merlo et al., 2006; Reiter et al., 2017; Gillies et al., 2018)* with driver mutations (single nucleotide variants, copy number alterations, translocations) conferring a selective advantage. Yet tumors differ in which driver mutations they carry *(Curtis et al., 2012; Ciriello et al., 2013)*. Understanding tumor diversity, comprehending the function of driver mutations in different contexts, and understanding why drugs affect some tumors and not others, are all pressing fundamental questions *(Vogelstein et al., 2013)*.

The growth in molecular data on tumors has driven the development of powerful algorithms to sort tumors into classes and clusters (Ronglai Shen, Olshen and Ladanyi, 2009; Dawson *et al*., 2013; Wang *et al*., 2017). These algorithms attempt to maximally separate tumors and cluster them according to molecular criteria. Cluster membership can be correlated with drug response and prognostics to help guide and design treatment for individual patients based on their molecular profile (Barretina *et al*., 2012; Heiser *et al*., 2012; Ciriello *et al*., 2013; Pemovska *et al*., 2013; Iorio *et al*., 2016).

While the ability to sort tumors is powerful and useful, there remains an open question of understanding, from a theoretical basis, *why* tumors vary in the way that they do. To address this, we apply a theory of multi-task evolution to the case of cancer. We reasoned that multi-task evolution may apply to cancer because cancer is a case of intense evolution inside the body that can play out over years, with generation times of cells that can be on the order of days (Driessens *et al*., 2012; Gillies, Verduzco and Gatenby, 2012; Labi and Erlacher, 2015; Lan *et al*., 2017). Furthermore, for cancer to grow and survive, it needs to fulfill multiple tasks, including growth, stress resistance, interaction with the immune system and so forth (Hanahan and Weinberg, 2011). Each task needs a different profile of gene expression – ribosomes for growth and stress proteins for survival.

Presumably no tumor can be optimal at all tasks at once, because cells can only make a limited amount of protein per unit biomass, and proteins for different functions can interfere with each other. Thus, cell communities that optimally manage the tradeoff relevant for their particular niche in the body will outgrow and out-survive cells that are suboptimal.

Such tradeoffs are well-known in bacteria: cells that grow faster are more sensitive to stress and antibiotics (Balaban *et al*., 2004). Cells that grow in a challenging environment need to express survival genes, which comes at the expense of growth genes (Scott *et al*., 2010; Shoval *et al*., 2012). A similar tradeoff is found in cancer cells (Aktipis *et al*., 2013). For example, cancer cells exposed to hypoxia can survive by invading the tissue surrounding the tumor (Sullivan and Graham, 2007; Hatzikirou *et al*., 2012) which can come at the cost of a reduced proliferative activity (Evdokimova *et al*., 2009; Tsai and Yang, 2013).

But growth and survival are only two of the possible tasks that affect tumor cell fitness. How can we detect and understand tradeoffs between three and more tasks? How can we identify the tasks without assuming what the tasks are a-priori? Going beyond a tradeoff between two tasks requires special approaches that can detect the impact of multiple simultaneous tasks.

To address the question of tradeoffs in tumors between multiple tasks, we apply multi-task evolutionary theory, known as Pareto task inference (ParTI), to cancer. We use ParTI to (i) identify tradeoffs between five universal tasks shared across cancer types, (ii) show that tumors that specialize in a task are differentially sensitive to drugs that disrupt that task, and (iii) demonstrate that each driver mutation moves gene expression towards specific archetypes and hence towards specialization in specific tasks. This suggests a picture of tumor diversity based on multi-task evolution. Thus, our goal is not to separate tumors, as is already done well by existing algorithms, but to add to our understanding of which evolutionary tradeoffs lead to the observed variation between tumors, and to explain drug sensitivity and driver mutations in terms of the concept of specialist/generalist tumors at different tasks.

## Results

The starting point for ParTI is that a tumor needs to perform multiple tasks in order to thrive *(Hanahan and Weinberg, 2011)*, but that these tasks are not known a priori. ParTI further assumes that no tumor can be optimal at all tasks at once, leading to a fundamental tradeoff. Due to the intense competition and high turnover of cells in a tumor, natural selection is expected to be strong (Driessens *et al*., 2012; Gillies, Verduzco and Gatenby, 2012; Labi and Erlacher, 2015; Lan *et al*., 2017). Cells with sub-optimal gene expression will lose the competition to cells with gene expression that is more optimal given the tradeoffs. It thus makes sense to apply Pareto optimality theory to the tumor situation. The theory predicts that such tradeoffs lead to a characteristic pattern: gene expression, averaged over all of the cell types in the tumor, is arranged within a polygon or polyhedron in gene expression space, with flat sides and sharp vertices *(Shoval et al., 2012)*. The vertices of the polyhedron, called archetypes, are profiles optimal for one of the tasks (Fig. 1A). Specialist tumors at a task lie close to a vertex, and generalists lie in the middle of the polyhedron (Fig. 1B). Tumors outside the polyhedron are sub-optimal, and will not be selected.

**Fig. 1.**
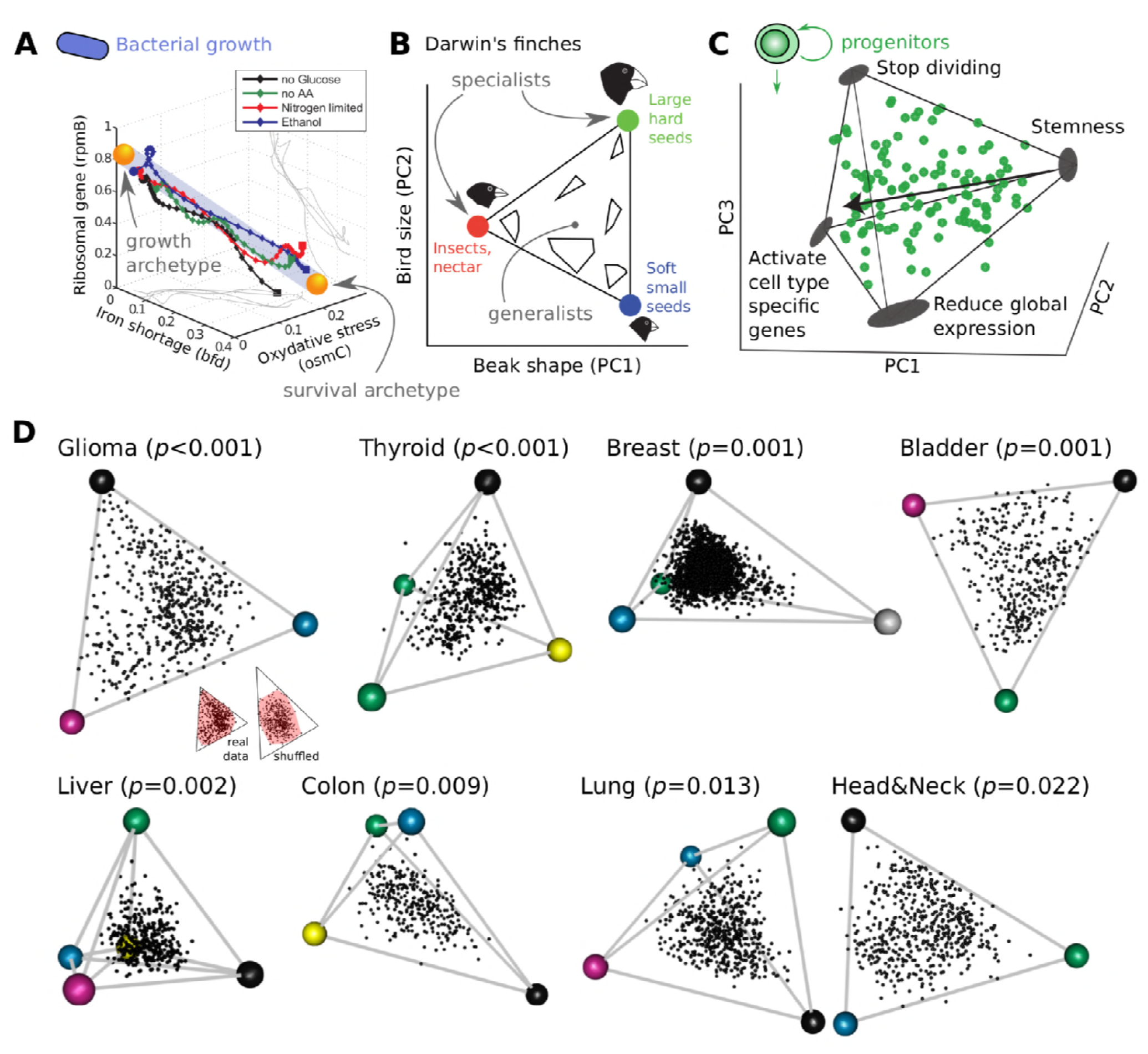
Tradeoff between tasks leads to phenotypes in a polyhedron, whose vertices are archetypes specializing in one task. **A.** 90% of the variation in *E. coli* gene expression falls on a line due to a trade-off between tasks of growth and survival. Axes are percent of total promoter activity. **B.** Morphology of Darwin’s ground-finch species falls on a triangle. Specialists in different diets are found near the three archetypes, and generalist are near the center of the triangle. A and B adapted from *(Shoval et al., 2012)*. **C.** Single-cell gene expression of mouse intestinal progenitor cells fall on a tetrahedron, shown in principal-components (PC) space. The four archetypal gene expression profiles correspond to fundamental progenitor cell tasks. Adapted from *(Korem et al., 2015)*. **D.** Tumor gene-expression profiles of 8 cancer types fall on polyhedra. Individual tumors (dots) plotted in the space spanned by the first three gene-expression PCs (TCGA, breast cancer from Metabric). Archetype (colored dots) number and position were inferred using ParTI. Inset: shuffled data has a convex hull (CH, pink) that fills less of the minimal enclosing (ME) triangle than the real data. The ratio of the CH area (volume) and ME triangle (or tetrahedron) was used to compute statistical significance.

Thus, finding polyhedral structure in data allows one to infer the number and nature of the tasks. Such polyhedral structures, tasks and tradeoffs were found in several contexts including bacterial and stem-cell gene expression and animal morphology (Fig. 1A-C) (Shoval *et al*., 2012; Korem *et al*., 2015) and in a preliminary analysis of breast cancer (Hart *et al*., 2015).

To test whether human tumor transcriptomes fall on low-dimensional polyhedra, we analyzed the transcriptomes of primary tumor samples from TCGA *(McLendon et al., 2008)* and Metabric *(Curtis et al., 2012; Pereira et al., 2016)* (normal samples were removed). We used the Pareto Task Inference (ParTI) software package *(Hart et al., 2015)* which fits lines, triangles, tetrahedra and so on to data, finds the best fit polyhedron and calculates its statistical significance. ParTI thus infers the number of archetypes and their position in gene expression space. We tested the 15 cancer types that have at least 250 primary tumor samples. We find that polyhedra with 3-5 archetypes describe gene expression of 6 cancer types, including breast, colon, thyroid, bladder, low-grade glioma and liver (Fig. 1D, Fig. S1A, FDR<10%, p<0.001 to p=0.009), with two more showing borderline significance (lung, p=0.01 and head&neck, p=0.02).

The 7 other cancer types (including kidney renal clear cell carcinoma and ovarian cancer) appear as clouds in gene expression space without detectable vertices; possible reasons include having primarily one task and hence no strong tradeoffs, having too many tasks and thus too many vertices to resolve, or data heterogeneity not currently understood.

We find that the archetypes of the polyhedron for different cancer types are similar to each other in terms of gene expression (Fig. S1D-E). We therefore hypothesized that tumors from different cancer types face similar trade-offs. To test this, we pooled the 3180 primary tumors from the 6 cancer types, after correcting gene expression profiles for tissue identity (normalizing each sample by the mean expression profile of its cancer type, Methods). We find that transcriptomes of the tumors vary in a continuum inside a polyhedron bounded by five archetypes (p=0.002, Fig. 2A). Tumors from different cancer types are spread widely within the polyhedron (Fig. 2A, Fig. S2A), and are found close to 3-5 of the archetypes depending on the cancer type (Fig. 2B, Fig. S2A-C).

**Fig. 2.**
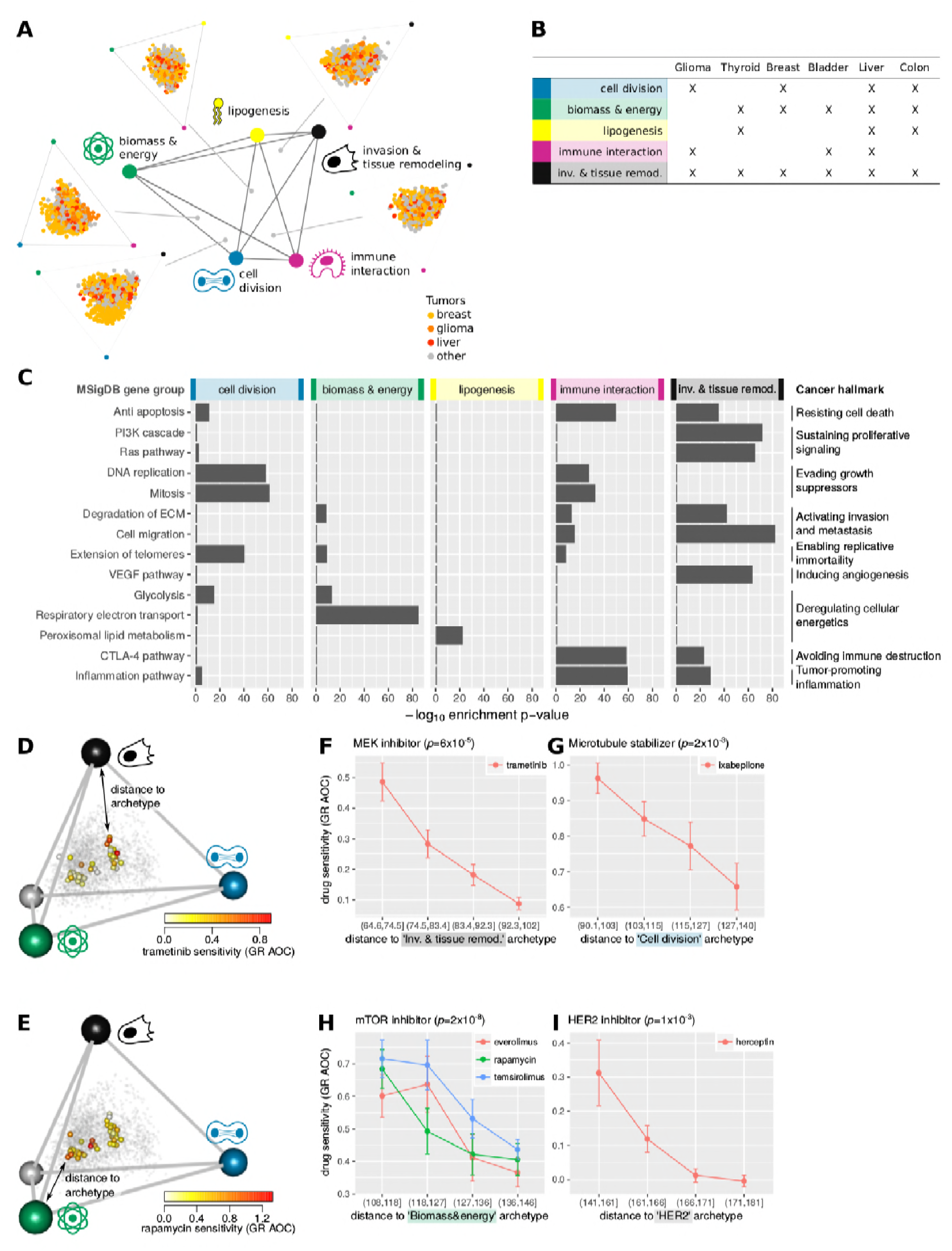
Tumors face trade-offs between 5 universal cancer tasks. **A.** Gene expression profiles of tumors from six cancer types fall in a continuum bounded by five archetypes, with indicated tasks. Tumors are shown projected on polyhedron faces (red-breast, blue-colon. gray-other tissue). Archetype number and position (colored dots) were inferred using ParTI. **B.** Universal cancer tasks found in each cancer type. X: significant overlap between cancer archetype and universal archetype in the set of genes whose expression is enriched in tumors closest to archetype (Methods). **C.** The five cancer tasks represent hallmarks of cancer *(Hanahan and Weinberg, 2011)*. For each cancer hallmark, one to three representative MSigDB gene groups (GG) were chosen. P-value are for enrichment in the 5% of the tumors closest to each cancer archetype. For visualization, p-values for GG that are not significant at FDR<0.1 or are not maximally enriched at a given archetype are set to 1. **D-E.** Breast cancer (BC) cell-line gene expression *(Heiser et al., 2012)*, projected on the BC tumor tetrahedron. Color: sensitivity to indicated drug. **F-I.** Sensitivity to indicated drug, defined as area under curve (AOC) of growth rate (GR) dose response, as function of Euclidean distance of cell-line gene expression to indicated archetype.

To infer the tasks performed by these five universal archetypes, we analyzed which pathways and functional gene groups are expressed highest in the tumors closest to a given archetype, using MSigDB *(Subramanian et al., 2005)* (Database S1, FDR<10%, using leave-one-out controls). We also determined which clinical properties are frequent among the tumors closest to a given archetype compared to other tumors in the dataset (Database S2-S3, FDR<10%).

We find clear tasks for each of the five archetypes (Table 1, Fig. S2D). The five tasks are: cell division, biomass & energy production, lipogenesis, immune interaction and invasion & tissue remodeling. The tasks match the hallmarks of cancer defined by *(Hanahan and Weinberg, 2011)* (Fig. 2C). A given hallmark can contribute to one or more tasks.

Tumors from patients with higher number of invaded lymph nodes are found near the invasion & tissue remodeling archetype (p<10^−3^, Database S3). Tumors with highest histological grade (which corresponds to poor tissue differentiation) are found near the immune interaction archetype (p=10^−6^, Database S2). Tumors at an earlier stage (Stage II) are found near the cell division, biomass & energy and lipogenesis archetypes (p<0.003) whereas late tumors (Stage III) are found near the immune interaction and invasion & tissue remodeling archetypes (p<10^−5^, Database S2).

The tasks for each tissue type are indicated in Fig. 2B, and color coded on the archetypes of Fig. 1B. Each tissue type seems to show a trade-off between 3-5 universal tasks (Fig. 2B, Fig. S2C). Breast cancer shows three universal archetypes (division, invasion, biomass & energy) and a fourth specific one enriched with HER2-positive tumors (p<10^−8^, Database S2), which seems to be tissue specific (Fig. S2C).

### The polyhedra do not result from averaging over different cell types

In interpreting the task of the archetypes, one concern is that we use data averaged over all cells in the tumor. The different archetypes could represent individual cell types (immune cells, stromal cells, malignant cells, …) rather than tasks. Such a situation would also result in polyhedra: if tumors are weighted average of cell types, they fall on a polyhedron with pure cell types at the vertices.

However, the data is inconsistent with the hypothesis that archetypes represent individual cell types. If archetypes represented individual cell types, tumors should fall on polyhedra in *linear* gene expression space. We find no significant polyhedra in linear gene expression space, only in log gene expression space (Fig. S1A-B). Furthermore, the cell division, biomass&energy and lipogenesis archetypes show high purity of cancer cells(Fig. S1C). These observations suggest that archetypes do not represent specific cell types.

### Specialist tumors are sensitive to drugs that interfere with their task

To further test the hypothesis that tumors face trade-offs between conflicting tasks, we reasoned that tumors whose gene expression is near an archetype are task specialists, and hence will be most sensitive to drugs which specifically disrupt that task. To test this hypothesis, we used data from Heiser et al. who assessed the sensitivity of 49 human breast cancer cell lines to a panel of 77 drugs *(Heiser et al., 2012)*. This dataset includes growth rate, overcoming a limitation of larger datasets in which apparent drug sensitivities cannot be corrected for growth rate effects *(Hafner et al., 2016)*.

To determine the position of these cell lines relative to the breast cancer archetypes, we projected the transcriptome of the cell lines onto the geneexpression space defined by breast tumors (Fig. 2D-E, Methods).

We find that cell-lines closest to the invasion and tissue remodeling archetype are sensitive to trametinib, an inhibitor of the Ras pathway is up-regulated in tumors close to the invasion and tissue remodeling archetype (Fig. 2F, Table 1). Similarly, cell lines closest to the cell-division archetype are sensitive to ixabepilone which stabilizes microtubules, and thus targets mitosis (Fig. 2G). Cell lines near the biomass&energy archetype are most sensitive to drugs which inhibit mTOR (Fig. 2H), a controller of cell growth *(Laplante and Sabatini, 2012)*. Finally, cell lines close to the breast-cancer-specific HER2 archetype (tumors that over-expresses the erbB-2 receptor) are sensitive to herceptin, an erbB-2 inhibitor (Fig. 2I). This differential sensitivity to drugs supports the hypothesis that tumors close to archetypes are task specialists.

### Driver mutations push gene expression to specialize in specific tasks

Finally, we asked how genetic alterations in tumors fit into the present trade-off picture. We computed the mean effect of each genetic alteration, a vector that describes how this alteration shifts gene expression (the difference in gene expression between tumors with and without the alteration) (schematically shown in Fig. 3A). We compared this effect vector with the polyhedron for each cancer type. There are two possible situations: the effect vector can align with the polyhedron, or instead can point away from the polyhedron. To visualize this, if the front were a triangle on a plane, the effect vector could lie on the same plane (have a small angle with the plane), or could point away from the plane (have a large angle) (Fig 3A). Importantly, shuffled controls typically point away, (angle=60°-80°) because the polyhedron explains only a fraction (20-40%) of the variation in the data.

**Fig. 3.**
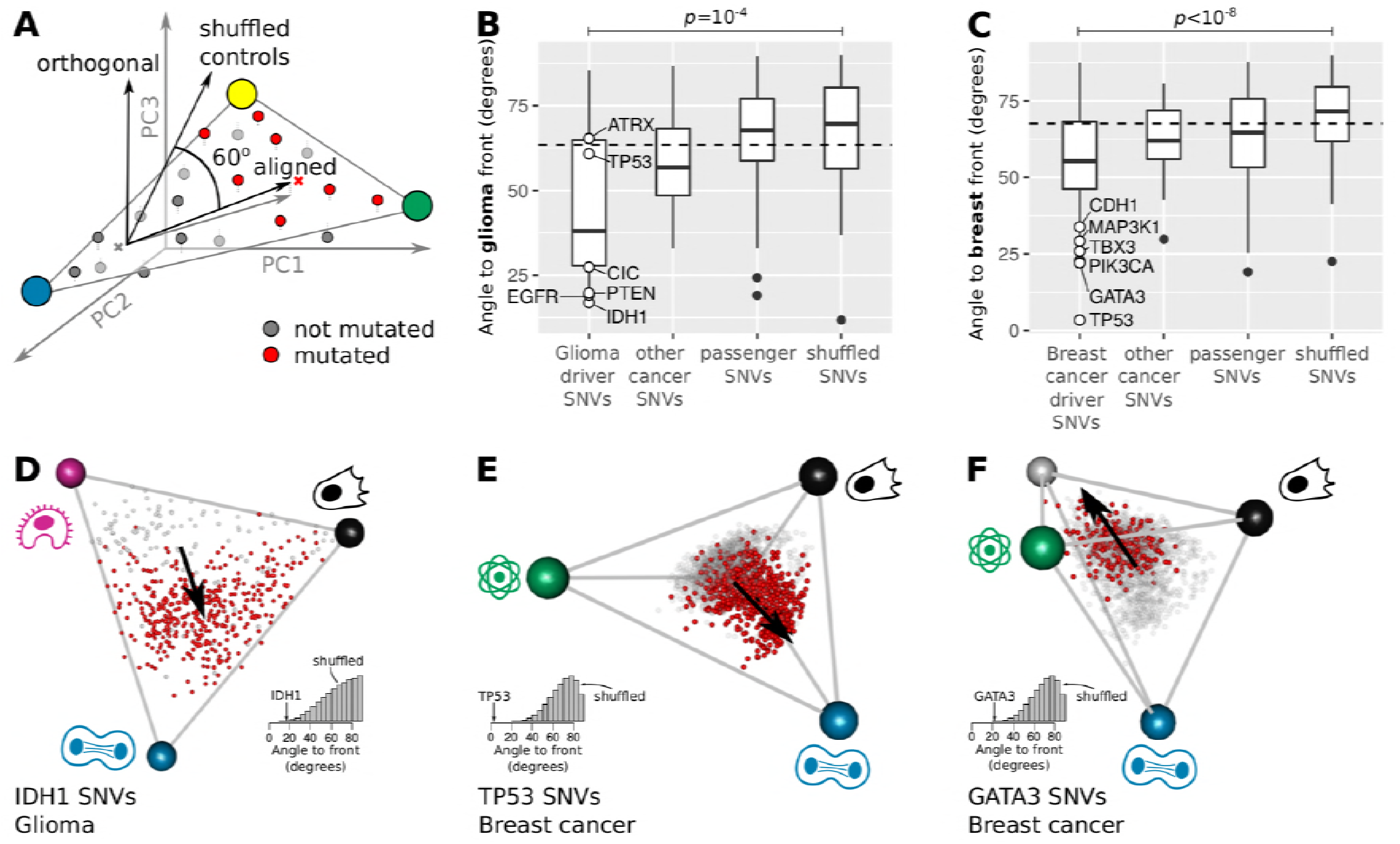
Driver mutations push tumor expression towards specialization in specific tasks. **A.** The effect of a mutation on gene expression is defined as the vector connecting the centroids of tumors without (grey) and with (red) the mutation (schematic). Alignment of mutation effect vector to the front is defined by the angle of the effect vector to the sub-space defined by the front. Shuffled controls show a 62°-75° angle with the front depending on cancer type. **B-C.** In glioma (B) and breast cancer (C), driver mutations are better aligned to the polyhedron formed by the archetypes than passenger mutations and random mutations. White dots: 6 most frequent driver SNVs in that cancer type. **D.** In low-grade glioma, *IDH1* SNVs point towards the cell division archetype. **E.** In breast cancer, *TP53* SNVs point towards the cell division archetype. **F.** In breast cancer, *GATA3* SNVs point towards the face defined by the lipogenesis, invasion & tissue remodeling and HER2 archetype.

Strikingly, for five cancer types, the effect vectors of driver mutations align with the polyhedron much more closely than expected from shuffled data: glioma (p=10^−4^, Fig. 3B), thyroid cancer (p=10^−3^), breast cancer (p<10^−8^, Fig. 3C), bladder (p=0.02), colon cancer (p=5×10^−3^, Fig. S3A). Drivers are also much more aligned than non-driver cancer genes and passenger mutations – collected from *(Santarius et al., 2010; Ciriello et al., 2013; Gonzalez-Perez et al., 2013; Nik-Zainal et al., 2016; Pereira et al., 2016)* - the latter are as aligned as shuffled data (Fig. S3A).

We find that driver mutations move gene expression towards specific archetypes. For example, *IDH1*, a strong driver in glioma, shifts gene expression towards the cell-division archetype (Fig. 3D). In breast cancer, the common *TP53* mutation is the most aligned with the front. It points directly towards one archetype, cell division (angle to archetype = 18°, mutation enriched 2.6-fold in the 5% of tumor closest to archetype, p<10^−16^). Mutations in *TP53* in breast cancer and *IDH1* in glioma thus coordinate gene expression towards specializing in the cell-division task. Another breast cancer driver, *GATA3*, shifts gene expression towards the face defined by the lipogenic, *HER2* and invasion & tissue remodeling tasks and away from the cell division archetype (p=0.003). The same conclusion is found for all 21 drivers with significant alignment to the polyhedra (Database S4 lists the driver mutations and the archetypes they point to, FDR<10%). Thus, aligned driver mutations can be interpreted as knobs that tune gene expression towards some tasks and away from others. Although the tasks are universal, the drivers that shift gene expression toward each task are often tissue specific.

We also analyzed copy number alterations (CNAs). CNAs show the same features found for SNVs above: driver CNAs have effect vectors that are highly aligned with the polyhedron (Fig. S3B) and push gene expression to specialize in specific tasks (Fig. S3C). For example, *PTEN* deletion in lower grade glioma points to the immune archetype and *MYC* amplification in breast cancer points at the cell division archetype (inferred tasks for 229 CNAs are listed in Database S5, FDR<10%).

## Discussion

The goal of this study is to provide a framework to understand the diversity of tumors in terms of multi-task evolution. We used ParTi to discover the number of tasks and their biological nature. The present work suggests trade-offs between 5 tasks. While the proliferation vs survival tradeoffs was already documented in cancer, our results suggest that there are other tradeoffs that could be of comparable importance in shaping tumor gene expression. We suggest tradeoffs between organizing metabolism to grow on glucose vs using lipids for growth, and between rapid cell division, immune evasion and tissue remodeling. Testing these tradeoffs could be the object of follow-up experiments aimed at measuring performance at the five tasks by quantifying DNA replication rate, lipid metabolism, protein synthesis rate, immuno-resistance and invasion rate in different tumors. If tradeoffs are at play, one should find negative correlation between the different measures of performance.

This framework predicts that tumors that specialize in a task should be more sensitive to drugs that impair that task. We find evidence for this prediction (Fig. 2D-I). Future studies on specific tumors with more samples and accuracy can probably uncover additional archetypes. These can offer hypothesis for which drugs to use on which tumor, and which combinations might work for generalists that tradeoff between multiple tasks.

Furthermore, we find that most diver mutations tune gene expression not in arbitrary directions in gene-expression space, but instead towards specialization in specific tasks. Non-driver mutations have more arbitrary effects on gene expression, and do not point towards specific archetypes. This could provide an approach to detect driver mutations and differentiate them from non-driver mutations.

Note that our goal is not to discover new tasks or cancer subtypes: existing approaches to sort tumors are better for this task (R. Shen, Olshen and Ladanyi, 2009; Dawson *et al*., 2013; Wang *et al*., 2017). Such sorting approaches are typically not based on a theoretical evolutionary basis, but instead aim to maximize the difference between tumors according to molecular features. The point of the present study is not to offer an improved way to sort tumors, but to provide a unifying explanatory framework to rationalize why tumors vary in the way that they do based on multi-task evolution. From this point of view, it is gratifying that most of the tasks we find are well-known, and the sensitivity to drugs is easily understandable in terms of known mechanisms.

We considered here total gene expression in tumors. Future work can test the present framework at the level of single-cell variation inside a tumor (Korem *et al*., 2015; Tirosh *et al*., 2016; Kim *et al*., 2018). One prediction, based on concepts from ecology (Schluter, 1996; Sheftel *et al*., 2018) is called “evolution along lines of least genetic resistance”-the finding that the main axes of variations between individuals in a given species is aligned with variation between species in the same taxon. This predicts, in the case of cancer, that variation between single cells in a tumor will align with variation between different tumors, because of the shared tasks (Shoval *et al*., 2012).

In summary, we suggest a framework for understanding tumor variation based on evolution under tradeoffs between tasks. Tumor gene expression lies in a continuum in a polyhedron whose five vertices are archetypal expression programs for five tasks that recur in different cancer types. Tumors can be specialists at a task or generalists: specialists have gene expression close to a vertex and generalists lie in the middle of the polyhedron. We find support for the hypothesis that specialists in a task are more sensitive to drugs that disrupt that task. Driver mutations are often like knobs that tune gene expression towards specialization in specific tasks. This framework, if validated by further research, offers a way to understand tumor variation in terms of task specialization.

## Acknowledgments

The authors would like to thank Ruthie Shouval, Ravid Straussman and members of the Alon lab for discussions. We thank Oscar Rueda for discussions and assistance in obtaining data.

## Authors contributions

Conceptualization: J.H., C.C., U.A.; Methodology: J.H., P.S., C.C., U.A.; Software, formal analysis & investigation: J.H., P.S., N.B., A.Z., H.S.; Writing: J.H., C.C., U.A.; Supervision: J.H., U.A.; Funding Acquisition: J.H., C.C., U.A.

## Data and material availability

The results shown here are based upon data generated by the TCGA Research Network: http://cancergenome.nih.gov/. All data used in this paper and scripts written to analyze the data are available at in the supplementary material.

## Methods

### Data and software availability

Input data files as well as scripts to reproduce the analyses can be downloaded at: https://data.mendeley.com/datasets/2r9h9xzwm3/draft?a=8dd0167f-fc1c-463b-8594-28e14ff094a7

### Data sources

- TCGA data (release 2015-02-24)

∘ Gene expression We downloaded the RSEM-normalized HiSeqV2 gene expression data (log2 RPKMs) from the TCGA data portal. The TCGA data portal is now retired but the data can be retrieved from the Genomics Data Commons portal of the National Cancer Institute. The starting point of our analyses were the ‘genomicMatrix’ files which contain expression levels for 20530 human genes in 15 cancer types. We considered all cancer types with at least 250 primary tumor samples:

▪ bladder (TCGA disease code: BLCA) (Weinstein et al. 2014)
▪ breast (TCGA disease code: BRCA) (Network 2012)
▪ cervical (TCGA disease code: CESC) (Burk et al. 2017),
▪ colon (TCGA disease code: COAD) (Muzny et al. 2012)
▪ head&neck (TCGA disease code: HNSC) (Lawrence et al. 2015)
▪ kidney (TCGA disease code: KIRC) (Creighton et al. 2013)
▪ lower grade glioma (TCGA disease code: LGG) (Network 2015)
▪ liver (TCGA disease code: LIHC) (Network et al. 2017)
▪ lung adenocarcinoma (TCGA disease code: LUAD) (Collisson et al. 2014)
▪ lung squamous cell carcinoma (TCGA disease code: LUSC) (Hammerman et al. 2012)
▪ ovarian cancer (TCGA disease code: OV) (Bell et al. 2011)
▪ prostate (TCGA disease code: PRAD) (Network et al. 2015)
▪ stomach (TCGA disease code: STAD) (Bass et al. 2014)
▪ thyroid (TCGA disease code: THCA) (Network et al. 2014)
▪ uterus (TCGA disease code: UCEC) (Getz et al. 2013)
∘ SNVs and CNAs From the TCGA data portal, we downloaded single nucleotide variant (SNV) calls, and copy number alteration (CNA) calls (gistic2 thresholded), as reported in ‘genomicMatrix’ files. For SNVs, we focused on genes mutated in at least 1% of the samples, to reduce the computational time and the memory requirements of our analyses. We also analyzed the CNAs which are prevalent. Since unlike mutations which can be present or absent, CNAs have 5 different values, we used an entropy measure, and analyzed the 1% of genes with highest entropy in their CNAs. For each gene, we determine the fraction of samples with 1. strong deletions f_1_, 2. weak deletion f_2_, 3. no detectable copy number alteration f_3_, 4. weak amplifications f_4_, and 5. strong amplification f_5_. We then computed the entropy of the corresponding distribution, -*Σ_i_ f_i_* log *f_i_*. A gene whose copy number is never altered has entropy 0. Genes with highest entropy show the most frequent copy number alterations.
∘ Clinical data From the TCGA data portal, we also downloaded the clinical data reported in files named ‘clinical_data’. Discrete and continuous clinical features require different statistical treatment in order to determine which clinical features are over-represented among tumors close to individual archetypes. We thus separated clinical features into discrete and continuous features by manual examination.
- Metabric data

∘ Gene expression & CNAs We downloaded the normalized microarray gene expression data (data_expression.txt), CNAs data (data_CNA.txt) and clinical data (data_clinical.txt) for the 1970 tumors of the Metabric cohort (Curtis et al. 2012) from the cBio portal (brca_metabric.tar.gz). We manually separated clinical features into discrete and continuous features, as we did for the TCGA data.
∘ To reduce computational time and memory requirements, we focused our analysis of CNAs on the 1000 genes whose CNAs were most significantly associated to changes in the mRNA abundance of that gene - taken from Table S30 of (Curtis et al. 2012) - supplemented with 124 known breast cancer driver genes (Nik-Zainal et al. 2016; Pereira et al. 2016; Santarius et al. 2010). We supplemented these genes with 1000 random genes not previously reported as drivers, which we used as controls.
∘ We downloaded SNVs in 173 genes from (Pereira et al. 2016) (somaticMutations.txt) together with the ‘tumorIdMap.txt’ file which maps the tumor IDs used in Pereira et al. to those used in Metabric. We used this mapping to convert all tumor IDs to Metabric IDs.

### Gene expression analysis

For each cancer type, we focused our analyses on primary tumors (field ‘sample_type’ set to ‘Primary Tumor’ in the TCGA clinical annotation), thus removing normal control samples as well as local and distant metastases. Doing so excludes the possibility that archetypes correspond to differences in metastatic host tissues or in disease state (healthy vs cancer).

We started from a matrix of samples *times* genes (samples x genes). Entries of the matrix represent log2 normalized RPKMs (see Data Source). The goal of the analysis was not contrast highly expressed genes to low expressed genes in a given tumor but to identify the main changes in gene expression across tumors. To identify these changes in gene expression, we subtracted the average expression (averaging over samples) from each gene. As a result of this transformation, entries in the samples x genes matrix represented log2 fold change in expression of a given gene in a given sample compared to the average expression of that gene in that cancer type. We performed principal component analysis (PCA) on the transformed samples x genes matrix. We did not scale log2 fold changes by the standard deviation prior to PCA. As a result, large and correlated changes in gene regulation affected principal components more than uncorrelated, small changes which are prone to measurement error.

### Fitting polyhedra to tumor gene expression data with ParTI

To find polyhedra in the gene expression data from each tumor, we used the ParTI (Pareto Task Inference) matlab software package (Hart et al. 2015). Briefly, the input to ParTI is a large-scale dataset such as a matrix of sample x gene expression. ParTI determines the position of archetypes in gene expression space. These archetypes define a polyhedron. How well a polyhedron fits the data is quantified by the ratio of the volume of the bestfitting polyhedron to the volume of the convex hull of the data (t-ratio) (Shoval et al. 2012). The t-ratio is always larger than 1 and approaches 1 when the data falls on a polyhedron (see Fig. 1D, glioma panel). ParTI then computes a p-value for the statistical significance of the polyhedron by re-computing the t-ratio on 1000 shuffles that conserve the distribution of loadings on each PC but not the correlation between the PCs (Hart et al. 2015).

To choose the number of archetypes, we attempted to fit 3, 4 or 5 archetypes to each cancer type. We chose the smallest number of archetypes that produced a statistically significant polyhedron (p<0.01). We did not attempt to find 6 or more archetypes because of the limited number of tumor samples.

### Clustering archetypes from different cancer types

We performed clustering analysis on the gene expression profiles of archetypes from the 6 cancer types with significant polyhedra (glioma, thyroid, breast, bladder, liver and colon) to determine if different cancer types share similar tasks. We first determined the gene expression profile of each archetype as the average gene expression profile of tumors closest to the archetype (i.e. tumors in the first 1^st^ distance bin, see “Gene and clinical enrichment analysis” section).

Since different tissues can express different genes, we compared archetypes by collapsing the expression of 20530 genes onto MSigDB pathways (Subramanian et al. 2005). We defined the regulation of a given MSigDB pathway as the average regulation of the genes in that pathway. Averaging gene expression from single genes into MSigDB pathways transforms the archetypes x genes matrix into a matrix of archetypes x pathways.

We focused our comparison of archetypes on MSigDB pathways significantly up-regulated in tumors close to at least one archetype in all 6 cancer types (FDR<10%, see “Gene and clinical enrichment analysis”).

One difficulty with visualizing which pathways are up-regulated at different archetypes is that the expression of certain pathways varied strongly across tumors (e.g. immune pathways) whereas variations in the expression of other pathways were more moderate (e.g. peroxisome lipid metabolism). To visualize what pathways are up-regulated at each archetype on a common color-scale, we scaled the expression of each pathway by its standard deviation across archetypes. To overcome the challenge of visualizing and interpreting hundreds of pathways on the same figure, we selected 38 pathways enriched in all tissue types and representative of each of the 10 hallmarks of cancer (Hanahan & Weinberg 2011) (see rows of Fig. S1D).

We clustered the archetypes from the different cancer types by Gaussian mixture modeling (mclust R package). The Bayesian Information Criteria suggested 5 mixtures. Archetypes clustered by tasks, not by cancer type: archetypes from a given cancer types are assigned to different Gaussian mixtures, each of which groups together archetypes from different cancer types. This observation suggests that the same tasks are relevant for different cancer types.

### Finding universal cancer archetypes

To determine the cancer tasks shared by tumors from different cancer types, we performed ParTI on the gene expression profiles of all 3180 primary tumors from the 6 cancer types with significant polyhedra. Grouping gene expression profiles of primary tumors from the 6 cancer types combines two sources of variation: 1. differences in genes expression between tissues, 2. differences in gene expression between individual tumors of the same tissue.

In finding universal cancer types, differences in gene expression between tissues are of little interest. For example, it is not surprising for tumors from a fatty tissue like breast to show higher expression of lipid metabolism genes while brain tumors show higher expression of neuronal genes. Instead, we are interested in which genes individual tumors up-regulate or repress compared to other tumors of the same tissue. To identify these changes, we subtract the mean expression (averaged over samples) from each gene prior to assembling the matrix of all samples x genes. As a result, the average expression of each gene is 0 within each tumor type, and thus across all tumors. Therefore, entries in the samples x genes matrix represent log2 fold change in the expression of a given gene in a given sample relative to the average expression of that gene in tumors from the corresponding cancer type.

TCGA tumors were collected using a different technology (RNAseq) and analysis pipeline than the metabric tumors (microarray). To ensure homogeneity in the gene expression data, we thus used the TCGA 1095 breast tumors instead of the 1970 metabric breast tumors.

We applied ParTI as described in the section “Fitting polyhedra to tumor gene expression data with ParTI”. ParTI identified 5 archetypes, which define a polyhedron in 4 dimensions.

To visualize the position of the tumors in this 4D space, we projected tumors on the 2D faces of the polyhedron. Each face is defined by three archetypes.

To project tumors on faces, we computed the two vectors connecting the first archetype to the two other archetypes. These two vectors define a linear basis for the face, which we orthogonalized using the Gram-Schmidt algorithm and normalized so each basis vector had norm 1. Multiplying the orthonormal matrix by a matrix of the 4D coordinates of all tumors and of the three archetypes defining the face yielded their projections on the face.

Finally, to exclude the possibility that tasks correspond to specific cancer types and confirm that tumors from individual cancer types are instead found close to multiple archetypes, we computed the fraction of tumors from individual cancer types found among the 10% tumors closest to each archetype (Fig. S2B). If tumors from all cancer types were evenly represented close to all archetypes, we would expect these fractions to be 10%. We observe that tumors from all 6 cancer types make up ~10% of multiple archetypes. This observation confirms that individual tasks are relevant to multiple cancer types.

### Matching archetypes from different tissues to universal cancer archetypes

To determine whether tasks found in individual cancer types matched the five universal cancer tasks, we compared MSigDB pathways up-regulated at each tissue-specific archetype with MSigDB pathways up-regulated at each universal archetype.

We focused the comparison on MSigDB pathways significantly up-regulated at the archetype (FDR<10%) and with log2 fold change larger than 0.1 to discard pathways with minor regulation. For each pair of tissue-specific and universal archetype, we asked how many pathways are up-regulated in both. We then tested if the number of pathways common to both archetypes was significantly higher than expected under the null hypothesis of random sampling from the union of pathways found at any archetype of that cancer type (hypergeometric test). We concluded that two archetypes were statistically similar when the p-value of the hypergeometric test was below 1% after Bonferroni correction. Results of this comparison are shown on Fig. S2C. For visualization, the p-value of non-significant comparisons was set to 1.

The task of cancer-specific archetypes was assigned to that of the most similar universal archetype (grey dots on Fig. S2C), provided that the similarity between the two archetypes was statistically significant.

Using this procedure, we found that each universal cancer task was assigned to at most one archetype within each cancer type. This is expected if tumors from different cancer types share the same tasks. One exception was found in thyroid cancer (THCA): both archetypes 1 and 3 matched the task of biomass and energy best.

Finally, archetype 4 of the metabric breast tumors (BRCA 4, which is associated to the HER2 subtype) matched none of the universal archetypes. One possible interpretation is that this archetype performs a breast cancer specific task enabled by over-activation of HER2 signaling.

### Enrichment analysis of clinical features, MSigDB pathways and individual genes

Having inferred the number of archetypes and their position in gene expression space using ParTI, we characterized the task of each archetype.

We did so using the methodology previously described (Hart et al. 2015). Briefly, we considered the 50 tumors closest to each archetype. Since there were at least 250 primary tumors in each cancer type, 50 tumors correspond to at most 20% of tumors. In cases where 50 tumors represented less than 5% of all tumors, we selected the 5% tumors closest to the archetype. In these tumors, we then searched for over-represented clinical features (Database S2, Database S3) and up-regulated MSigDB pathways (Database S1). We performed enrichment analysis in each cancer type (using cancer-specific archetypes, Fig. 1D) as well as by grouping tumors from all 6 cancer types in one analysis (using the universal cancer archetypes, Fig. 2A). When analyzing tumors from all 6 cancer types together, we also looked for up-regulation of individual genes (Database S1).

For each continuous feature (e.g. expression of MSigDB pathways and individual genes, quantitative clinical features such as age, recurrence free survival, or tumor weight), we tested if the feature took significantly higher values in tumors closest to the archetype compared to other tumors (Mann-Whitney U test). For discrete features (e.g. the presence of an SNV or of a CNA, and qualitative clinical features such as the gender of the patient, the pathological stage, the molecular subtype of the tumor), we tested whether the feature was over-represented in tumors closest to individual archetypes using the hypergeometric test.

We controlled for the false discovery rates (FDR) using the Benjamini-Hochberg procedure. For each archetype, we report all features with FDR<10% and whose prevalence peaks in among tumors of the first distance bin of that archetype.

When analyzing tumors from all 6 cancer types, some clinical features are specific to a single cancer type. For example, PAM50Calls are only defined for breast cancer since PAM50 is molecular classification of breast tumors. Others features are relevant to multiple cancer types, such as the percentage of stromal cells in a tumor (% of stromal cells in a tumor). To distinguish features relevant to multiple cancer types from tissue-specific features (defined as features only defined in tumors of a single cancer type), tissue-specific features were tagged with the TCGA code of the cancer type in Database S2 and S3. For example: we renamed PAM50Call to BRCA.PAM50Call.

Testing for up-regulation of MSigDB pathways close to an archetype has the caveat of circular inference, as genes from a given pathway are used both to infer the position of the archetype and determine its task through MSigDB enrichment analysis. To address this caveat, we use a leave-one-out strategy. For each enriched MSigDB pathway, we remove genes belonging to this pathway and infer the position of archetypes again using the remaining genes. We then test if this pathway is still up-regulated in tumors close to the archetype. If not, we discard the pathway from the enrichment analysis.

In our analysis of 3180 tumors from all 6 cancer types, we applied this leave-out-one strategy to all MSigDB pathways with more than 100 genes. These pathways are more likely to influence the PCA and the position of the archetypes compared to pathways with less genes. We found that all MSigDB pathways identified in the original enrichment analysis were also enriched in the leave-one-out analysis. This observation suggests that archetypes are supported by large groups of genes belonging to diverse pathways, and that circularity is not a concern in the present results.

### Inferring tasks

To infer the task of each archetype, we used the same approach as in previous studies (Hart et al. 2015; Korem et al. 2015): we examined what MSigDB pathways were maximally up-regulated and what clinical features maximally over-represented in the 5% tumors closest to each archetype. We then used these MSigDB pathways and clinical features as clues to the task performed by the archetype. This section describes how up-regulated MSigDB pathways and over-represented clinical features support the tasks we inferred.

- Cell division (archetype 3 in Database S1-S3)

∘ Tumors close to this archetype up-regulate genes involved in the cell cycle (KEGG CELL CYCLE, p<1e-60). Up-regulated genes are involved in different phases of the cell cycle, such as the M phase (REGULATION OF MITOTIC CELL CYCLE, p<1e-20) and the S phase (REACTOME DNA REPLICATION, p<1e-55). This suggests that cells from these tumors are not arrested at some point in the cell cycle but are instead dividing more than cells from other tumors. Increased cell division is consistent with the observed high cellularity in breast tumors close to this archetype (p<1e-6).
∘ Genes involved in extending telomeres are up-regulated in tumors close to this archetype (REACTOME EXTENSION OF TELOMERES, p<1e-40). Telomeres are repeated hexanucleotides that protect the extremities of chromosomes. In somatic cells, telomeres are shortened at each division, thereby acting as a division counter which limits how many times a cell can divide before it becomes senescent or dies (Blasco 2005). Cancer cells are thought to achieve replicative immortality by acquiring the capacity to extend their telomeres, which normal cells typically cannot. Up-regulation of genes involved in telomere extension is thus consistent with the task of cell division.
∘ Tumors close to this archetype up-regulate genes involved in maintaining chromosome integrity (REACTOME ACTIVATION OF ATR IN RESPONSE TO REPLICATION STRESS, p=1e-55; KEGG MISMATCH REPAIR, p=1e-35), perhaps as a strategy to remain viable despite DNA damage that accumulate over cell divisions. This strategy appears only partially successful: the median tumor close to this archetype harbors 750 more CNAs than the rest of tumors (p<1e-40).
∘ Clinically, early tumors are over-represented close to this archetype (pathologic stage: Stage IIA, p<1e-16). Among breast tumors, triple negative tumors are over-represented (BRCA.breast carcinoma estrogen receptor status: Negative, p<1e-16; BRCA.breast carcinoma progesterone receptor status: Negative, p<1e-16; BRCA.HER2 Final Status nature2012: Negative, p<1e-16). Patients carrying these tumors show a 334 days longer recurrence free survival than patients with other tumors on average (X_RFS, p<1e-5).
- Biomass & energy (archetype 2 in Database S1-S3)

∘ Tumors close to this archetype up-regulate ribosomal proteins (KEGG RIBOSOME, p<1e-70; MITOCHONDRIAL RIBOSOME, p=1e-90) and genes needed in translation (REACTOME PEPTIDE CHAIN ELONGATION, p=1e-77; REACTOME TRANSLATION, p<1-e73). This suggests increased protein synthesis and biomass production.
∘ Genes involved in the proteasome (KEGG PROTEASOME, p<1e-65) are also up-regulated. Increasing proteasome activity is thought to be a strategy used by cancer cells to cope with the increased translation of low-quality proteins due to misfolding-causing mutations and aberrant splicing (Deshaies 2014). Aberrantly spliced mRNAs can contain premature stop codons. Such premature stop codons are detected as aberrantly spliced mRNAs, and degraded through the mechanism of nonsense mediated decay (NMD) (Kervestin & Jacobson 2012). NMD is up-regulated in tumors close to archetype 2 (REACTOME NONSENSE MEDIATED DECAY ENHANCED BY THE EXON JUNCTION COMPLEX, p=1e-75). These observations are consistent with the view that upregulation of the proteasome helps tumors cope with a proteotoxic crisis.
∘ In addition, tumors close to archetype 2 up-regulate genes needed in respiration (REACTOME FORMATION OF ATP BY CHEMIOSMOTIC COUPLING, p<1e-80; REACTOME TCA CYCLE AND RESPIRATORY ELECTRON TRANSPORT, p<1e-76) and in glycolysis (BIOCARTA GLYCOLYSIS PATHWAY, p<1e-25). This increase in energy producing pathways may serve to support protein synthesis and biomass production which accounts for the majority of ATP consumed in growing cells (Lynch & Marinov 2015; Wagner 2005).
∘ Clinically, tumors close to this archetype are found in patients at an early cancer stage (pathologic T: T2, p<0.001).
- Lipogenesis archetype (archetype 1 in Database S1-S3)

∘ Up-regulation of genes involved at this archetype support the task of lipid metabolism reprogramming, an area of cancer research that is currently undergoing significant developments (Beloribi-Djefaflia et al. 2016).
∘ The main enzymes catalyzing de novo lipogenesis are up-regulated in tumors close to this archetype: ACLY (p<1e-11), ACACA (p<1e-16) and FASN (p<1e-7) (Menendez & Lupu 2007). Enzymes needed to synthesize glycosylphosphatidylinositols (GPIs) are also up-regulated in these tumors (KEGG GLYCOSYLPHOSPHATIDYLINOSITOL GPI ANCHOR BIOSYNTHESIS, p<1e-33). De novo lipogenesis has been proposed to promote growth and survival of cancer cells in several ways. First, de novo lipogenesis can support the need of cancer cells for membranes in cell proliferation (Menendez & Lupu 2007). Second, in the hypoxic environment of tumors cells, lack of oxygen blocks respiration which leads to an excess of reducing power (too much NADPH). De novo lipogenesis consumes NADPH and could thus help rebalance the redox balance (Menendez & Lupu 2007). Third, de novo lipogenesis produces lipids that are less sensitive to reactive oxidative species (ROS). ROS, which are produced by the mitochondrial activity that supports cell proliferation, can damage lipid membranes and thus endanger cell survival (Ward & Thompson 2012; Rysman et al. 2010). De novo synthesized lipids come saturated (Ackerman & Simon 2014). These saturated lipids are less sensitive to ROS than unsaturated lipids (Rysman et al. 2010). Subsequent desaturation requires oxygen, which is typically lacking in the tumor environment (Ackerman & Simon 2014). As a result, the increased levels of saturated lipids promoted by de novo lipogenesis may support cell survival.
∘ Peroxisomes are organelles most known for clearing reactive oxidative species. But they also carry out other functions, such as synthesizing lipids (in particular ether lipids) and shortening long fatty acid chains for use in mitochondrial metabolism (Lodhi & Semenkovich 2014; Wanders et al. 2015). Tumors close to archetype 1 up-regulate peroxisomal genes (peroxisomal part, p=1e-40) and genes involved in peroxisomal lipid metabolism (p<1e-22). Peroxisomal genes upregulated in tumors close to archetype 1 include ABCD3 (p=1e-22) which transports fatty acids to peroxisomes, as well as ACOX1 (p=1e-9), ACOX3 (p<1e-5) and AMACR (1e-17) which carry out beta-oxidation of long fatty acids in peroxisomes (Wanders et al. 2015). Although the exact function of peroxisomes in cancer is not clearly understood yet, beta-oxidation of long fatty acid chains in peroxisomes could represent an alternative energy source to glucose (Liu 2006).
∘ Clinically, tumors close to this archetype tend to be early stage (pathologic stage: Stage IIA, p<0.001) and well differentiated (neoplasm histologic grade: Low Grade, p<0.001). The median patient carrying these tumors was 8.5 years older than the median patient carrying other tumors (p<1e-4). Breast cancer tumors close to this archetype are enriched with hormonal cancer (BRCA.ER Status nature2012: Positive, p<1e-9, BRCA.PR Status nature2012: Positive, p<1e-8).
- Immune interaction (archetype 4 in Database S1-S3)

∘ Tumors close to this archetype up-regulate genes expressed in immune cells (KEGG ALLOGRAFT REJECTION, BIOCARTA TCYTOTOXIC PATHWAY) and related to immunity (BIOCARTA INFLAM PATHWAY, REACTOME INTERFERON GAMMA SIGNALING). There is also up-regulation of the PD-1 and CTLA-4 pathways which inhibit immune response (REACTOME PD1 SIGNALING, p<1e-55; BIOCARTA CTLA4 PATHWAY, p<1e-58). This suggests an archetype characterized by the invasion of tumors by immune cells, but whose action is inhibited. Consistent with the invasion of the tumors by immune cells, tumors close to this archetype show loss of tissue identity and poor differentiation in histological examinations (neoplasm histologic grade: High Grade, p=0.006).
∘ The median tumor close to this archetype has 5 more SNVs than other tumors (number of SNVs, p=0.0002). This association of the number of SNVs to immune invasion is consistent with previous reports that PD-1 blockage therapy is most effective against tumors with higher mutational burden (Rizvi et al. 2015).
∘ Clinically, patients with these tumors are at a more advanced disease stage (pathologic stage: Stage III, p<1e-5) than archetypes 1-3 (Stage II). This difference suggests that the task of immune interaction becomes relevant to tumors at a later stage than the tasks of cell division, biomass&energy and lipogenesis.
- Invasion and tissue remodeling (archetype 5 in Database S1-S3)

∘ Tumors closest to this archetype up-regulate genes involved in the remodeling of extracellular matrix (EXTRACELLULAR MATRIX STRUCTURAL CONSTITUENT, p<1e-65; EXTRACELLULAR STRUCTURE ORGANIZATION AND BIOGENESIS, p<1e-65; REACTOME DEGRADATION OF THE EXTRACELLULAR MATRIX, p<1e-40). Disorganization of the ECM participates in tumor progression and metastasis (Lu et al. 2012). The second most upregulated MSigDB pathways in these tumors is collagen (p=1e-57), which suggests the presence of cancer associated fibroblasts (CAFs). Accordingly, the average percentage of stromal cells in tumors close to this archetype was 11 points higher compared to other tumors (p=0.0006).
∘ Tumors close to this archetype up-regulate genes involved in invasion and metastases such as cell migration (p<1e-80), and angiogenesis (REGULATION OF ANGIOGENESIS, p<1e-64). Consistent with the task of invasion, tumors close to this archetype invaded 1.8 more lymph nodes than other tumors on average (number of lymphnodes positive by he, p<0.0005). Lymphatic and venous invasion is over-represented in colon tumors close to this archetype (p<0.0004), as well as lymphovascular invasion in bladder tumors (BLCA.lymphovascular invasion present: YES, p=0.003). Patients carrying tumors close to this archetype are at the most advanced disease stage of all 5 archetypes (pathologic stage: Stage IIIC, p<1e-7). Thus, both gene expression and clinical features support the task of invasion.
∘ In addition, tumors close to this archetype activate signaling pathways involved in development and tissue homeostasis: Hedgehog, Insulin-like growth factor, fibroblast growth factor, Wnt, TGFβ, Ras (PID HEDGEHOG 2PATHWAY, p<1e-80; REACTOME REGULATION OF INSULIN LIKE GROWTH FACTOR IGF, p<1e-65; REACTOME FGFR LIGAND BINDING AND ACTIVATION, p<1e-77; PID WNT SIGNALING PATHWAY, p<1e-61; KEGG TGF BETA SIGNALING PATHWAY, p<1e-75; RAS GTPASE ACTIVATOR ACTIVITY, p=1e-72). Tumors are thought to hijack these programs to support cancer progression (Hanahan & Weinberg 2011). In the case of TGFβ signaling for example, epithelial tumors hijack a program normally used in development and wound healing in which epithelial cells acquire the ability to invade into a tissue and resist apoptosis (Diepenbruck & Christofori 2016).
∘ These tumors also up-regulate immune pathways, although at lower levels than tumors close to the immune interaction archetype. In addition, the identity of upregulated immune pathways differs between archetypes 4 and 5. For example, tumors close to the invasion & tissue remodeling archetype up-regulate the lectin pathway (BIOCARTA LECTIN PATHWAY is the 4^th^ most upregulated MSigDB gene group, p<1e-59). In contrast, the lectin pathway is not upregulated among tumors close to the immune interaction archetype.
∘ Finally, tumors close to this archetype up-regulate neural pathways (REGULATION OF NEUROGENESIS, VOLTAGE GATED SODIUM CHANNEL ACTIVITY). Voltage-gated sodium channel genes define the single most up-regulated MSigDB pathway among tumors close to this archetype. Upregulation of these pathways could result from cancer cells hijacking pathways normally active during neuron migration (Hanahan & Weinberg 2011). It could also be due to perineural invasion, a process in which cancer cells spread along nerves (Liebig et al. 2009). Alternatively, this up-regulation of neural pathways could be the sign of neural stimulation or modulation of the tumor microenvironment (Venkatesh & Monje 2017).

### Matching cancer tasks to cancer hallmarks

To compare the 5 universal cancer tasks to the 10 cancer hallmarks defined by (Hanahan & Weinberg 2011), we picked 1-3 MSigDB pathways representative of each cancer hallmark (Fig. 2C). We then plotted the p-value quantifying the statistical significance of the up-regulation of each MSigDB pathway in the 5% tumors closest to each archetype, as described in the “Enrichment analysis” section.

### Drug sensitivity analysis in breast cancer cell lines

To test if cancer cells that specialize in a task are more sensitive to drugs that interfere with the task, we used the gene expression and drug sensitivity data in 56 breast cancer cell lines of (Heiser et al. 2012).

We determined the position of the 56 cancer cell lines relative to the four breast cancer archetypes. To do so, we only considered the 14844 genes whose expression was quantified in both the 1970 Metabric breast tumors(Curtis et al. 2012) and in the 56 breast cancer cells lines (Heiser et al. 2012). We then performed quantile normalization on the joint samples x genes matrix of log2 gene expression using Bioconductor’s normalize.quantile function (Gentleman et al. 2004). To focus the analysis on the diversity in gene expression within cancer cell lines, we subtracted the mean gene expression profile out of each cell line. As a result, each gene had an average expression of zero, so that expression of a given gene in a certain cell line represented log2 fold changes relative to the mean expression across all breast cancer cell lines. We then projected both cell lines and breast cancer archetypes on the first three principal components of breast tumors, which were determined while finding the archetypes of breast cancer (see section “Fitting polyhedra to tumor gene expression data with ParTI”). In computing the projection, we kept only the 14844 dimensions corresponding to genes quantified both in tumors and cell lines. For visual reference, the 1970 Metabric tumors were also projected onto the same space.

GR AOC is a measure of drug sensitivity robust to cell-to-cell variations in growth rates (Hafner et al. 2016). We determined how the GR AOC varied as a function of euclidean distance to the archetype by grouping cell lines in four distance bins. For each bin *i*, we computed the median GR AOC *G_i_* and the within-bin standard error *σ_i_*. Because some drugs were assayed against only some of the 56 cell lines, we discarded drugs with less than 5 cell lines per bin.

For each archetype, we looked for drugs whose potency peaked in cell lines closest to the archetype, and then monotonically decreases away from it. To address the concern that the small sample size (56 breast cancer cell lines vs 1970 breast tumors per cancer type) can produce false positive drug-archetype pairs, we used a more stringent criteria to identify drugs effective against individual archetypes than the criteria used to find overrepresented MSigDB pathway and clinical features (see “enrichment analysis” section). We scored each drug by the product of the decrease in GR AOC in consecutive bins plus twice the standard error within bin, ⊓_i_ (G_i_ – G_i+1_ + 2 <*σ_i_*>), with <*σ_i_*> the median standard error across all bins. Drugs which score high on this scheme see their GR AOC peak close to the archetype and then steadily decrease away from it. If GR AOC increased by more than twice the standard error in any consecutive bins, the score was set to 0. This tolerates small bin-to-bin increase in GR AOC which could be due to measurement error or cell-to-cell variability. Finally, we tested if GR AOC was significantly higher in cell lines of the first distance bin compared to all other bins (FDR<10%, Mann-Whitney test, see “enrichment analysis” section). In doing so, everolimus, rapaycin and temsirolimus were grouped together as they share the same mechanism (mTOR inhibition, Fig. 2H).

Of the top 10 scoring drugs, 6 appear on Fig. 2F-I. In addition to these 6 drugs, Triciribine which inhibits the Akt kinase (an mTOR activator) is effective against the biomass & energy archetype (p=0.01), consistent with the sensitivity of the biomass & energy archetype to mTOR inhibitors (Fig. 2H). Gemcitabine, another drug, was not statistically significant at FDR<10% (p=0.11). The identity of the remaining drugs was kept confidential by the authors of the dataset (Heiser et al. 2012), thereby preventing further interpretation.

### Obtaining driver genes, other cancer genes and passenger SNVs and CNAs

To test the hypothesis that driver SNVs and CNAs are knobs that tune tumor gene expression towards specific tasks, we first compiled lists of 1. drivers alterations, 2. passengers alterations, and 3. alterations commonly found in cancer although not thought to be drivers. Second, we quantified whether these alterations pushed tumor gene expression along the cancer front or away from it.

For each cancer type except breast cancer, we obtained lists of driver genes from the IntOGen database (Gonzalez-Perez et al. 2013). We focused on known driver genes, for which there is causal evidence of their implication in cancer according to the Cancer Gene Census (Futreal et al. 2004). SNVs and CNAs in these known driver genes were labeled *driver* alterations.

In breast cancer, instead of using the IntOGen database, we took advantage of extensive efforts in classifying driver SNVs and CNAs as oncogenes and tumor suppressors (Santarius et al. 2010; Pereira et al. 2016; Nik-Zainal et al. 2016). We defined driver alterations from:

1. genes lying within minimal amplification regions, up-regulated as a result of this amplification, and with significant experimental evidence of their involvement in cancer development by Santarius et al. (class I, II and III) (Santarius et al. 2010). We defined these genes as oncogenes.
2. genes with a high proportion of recurrent SNVs (for oncogenes) or a high proportion of inactivating mutations (for tumor suppressors) in the survey of 2433 primary breast tumors of Pereira et al. (Pereira et al. 2016).
3. genes commonly amplified, or carrying mis-sense or inframe mutations (for oncogenes), and genes commonly deleted or carrying frameshifting or nonsense mutations (for tumor suppressors) in a survey of 560 breast tumors (Nik-Zainal et al. 2016).

SNVs corresponding to genes identified as oncogenes or tumor suppressors in any of these three studies were defined as *driver SNVs*. Amplifications (weak or strong) corresponding to genes identified as oncogenes were defined as *driver CNAs*. Deletions (weak or strong) corresponding to genes identified as tumor suppressors were also defined as *driver CNAs*.

Individual SNVs and CNAs that did not concern known driver genes were called *other cancer genes* if they appeared in the list of genes frequently amplified or mutated in cancer, listed in Table S2 of (Ciriello et al. 2013). Finally, SNVs and CNAs in genes not found in the two lists were called *passengers*.

### Assessing alignment of SNVs and CNAs to front

We represented the average effect of each alteration on gene expression as a vector connecting the centroid of tumors having the alteration to the centroid of tumors without the alteration (Fig. 3A). To quantify whether an alteration pushes gene expression along the cancer front or away from it, we computed the angle between the alteration vector and the cancer front. The cancer front was defined as the linear subspace that contains the archetypes. The angle between the cancer front and the alteration vector was computed by projecting the alteration vector on the cancer front, and by computing the scalar product between the alteration vector *u* and its projection *v*. The scalar product <*u, v*> was converted to an angle *α* (in degrees) using the inverse cosine, *α = 180* cos^−1^(<*u, v*>/|*u*|.|*v*|)/*π*.

To quantify the statistical significance of the alignment of an alteration vector to the cancer front, we randomly assigned alterations to tumors and computed the corresponding alteration vectors. By repeating this procedure, we obtained 10^5^ *shuffled control* vectors and their angles to the cancer front. We used the distribution of angles of shuffled controls – which was independent of the frequency of the alteration – to estimate the p-value that individual alterations vector are aligned to the cancer front. In principle, the p-value can be estimated from the fraction of shuffled controls with a smaller angle to the cancer front than a given alteration. An issue with this approach is that small angles are rare among shuffle controls. Thus, the p-values of strongly aligned alterations is imprecise. Precise estimation of the p-value of strongly aligned alterations is important when correcting for multiple testing.

To better estimate the p-values, we first computed the empirical cumulative distribution of the angle distribution of shuffle controls *F(α). 1-F(α)* is the p-value of an alteration of angle *α*. Because the function log[*1-F(α*)] is much flatter than *F(α)*, we focused on approximating *log[1-F(α)]. log[1-F(α)]* was fitted to a 5^th^ order polynom. In fitting the polynom, we used only angles *α* supported by at least 10 shuffled controls. By extrapolating the polynom to small angles α, we computed the p-value for each alteration vector to be aligned to the cancer front. Finally, we controlled the false discovery rate using the Benjamini-Hochberg procedure (Benjamini & Hochberg 1995).

## Supplementary Text

### Archetypes do not represent single cell types

We tested the possibility that the polyhedra described by tumors could be caused by mixing different cell types (cancer cells, fibroblasts, immune cells, …) in varying proportions rather than evolutionary trade-offs. Two observations suggest that it is unlikely that archetypes correspond to cell types which are mixed in varying proportions in different tumors.

First, mixing cell types in different proportions should produce tumors that describe polyhedra in linear gene expression space (Shen-Orr et al. 2010), but not log gene expression space. We looked for polyhedra in linear gene expression space by exponentiating gene expression right before subtracting the mean gene expression from each gene (see “Fitting polyhedra to tumor gene expression data with ParTI” in Materials and Methods). No significant polyhedra were found in linear gene expression space in any of the 15 cancer types.

Second, if archetypes represent individual cell types mixed in different proportions in different tumors, one archetype should represent pure cancer cells while the other archetypes should represent other cell types present in the tumor (fibroblasts, immune cells, …). Thus, tumor purity should peak at one of the archetypes and monotonically decrease away from this archetype, as cancer cells are increasingly mixed with other cell types. We tested this prediction by analyzing tumor purity, defined as the fraction of cancer cells in a tumor. Purity can be inferred from bulk tumor gene expression profiles by algorithms such as ESTIMATE (Yoshihara et al. 2013). We find that purity peaks at several archetypes (cell division, biomass&energy) and is lowest close to the invasion & tissue remodeling archetype (Fig. S1C-D). This observation is not expected if tumors mix cancer cells with other cell types in varying proportions but is consistent with the increased proportion of stromal cells found in tumors close to the invasion & tissue remodeling archetype (Database S3).

### Alignment of passenger CNAs to the front and spatial dependencies

In the five cancer types where driver SNVs aligned with the cancer front better than shuffled controls and with the exception of bladder, shuffled controls were as aligned to the cancer front as passenger mutations (Fig. S3A). In contrast, passenger CNAs were more aligned with the cancer front than shuffled controls in all 6 cancer types (Fig. S3B).

This result is likely explained by the fact that chromosomal amplifications or deletions typically involve a portion of a chromosome that contains many genes. Thus many neighboring genes are amplified or deleted along with the driver gene. One example of this phenomenon is shown on the right: the driver CNA ATM(-1) (i.e. ATM deletion) is best aligned with the cancer front, and pulls neighboring genes in its wake.

Because of the spatial dependencies of CNAs, the general pattern is that *driver* CNAS are most aligned with the front, followed by CNAs of *other cancer genes*, followed by *passenger* CNAs, followed by *shuffled controls* (Fig. S3B).

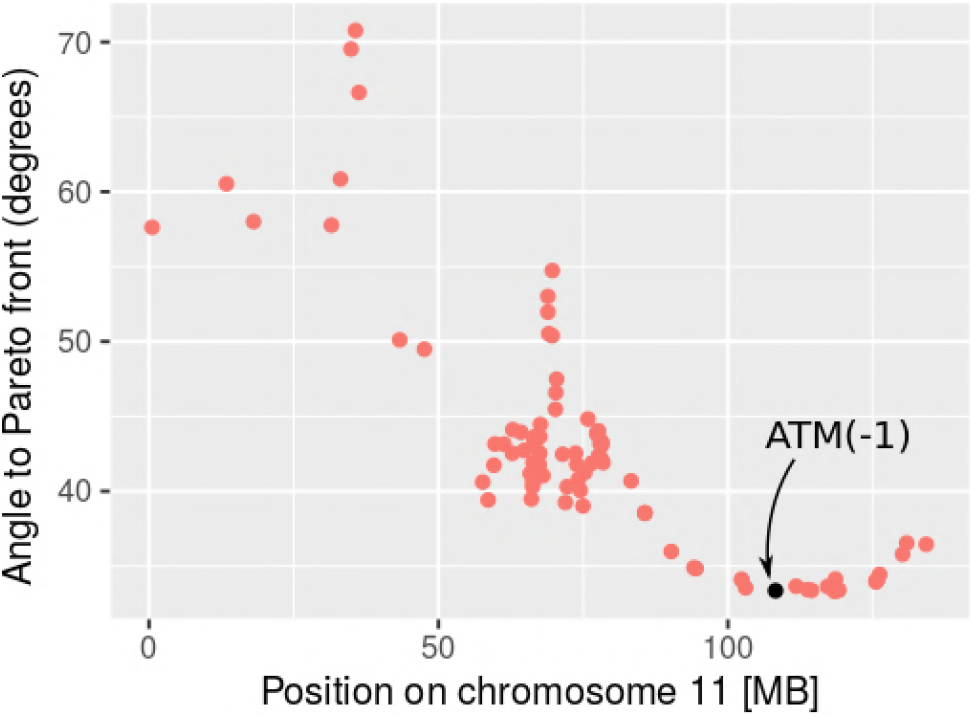

## Supplementary figures and tables

**Fig. S1.**
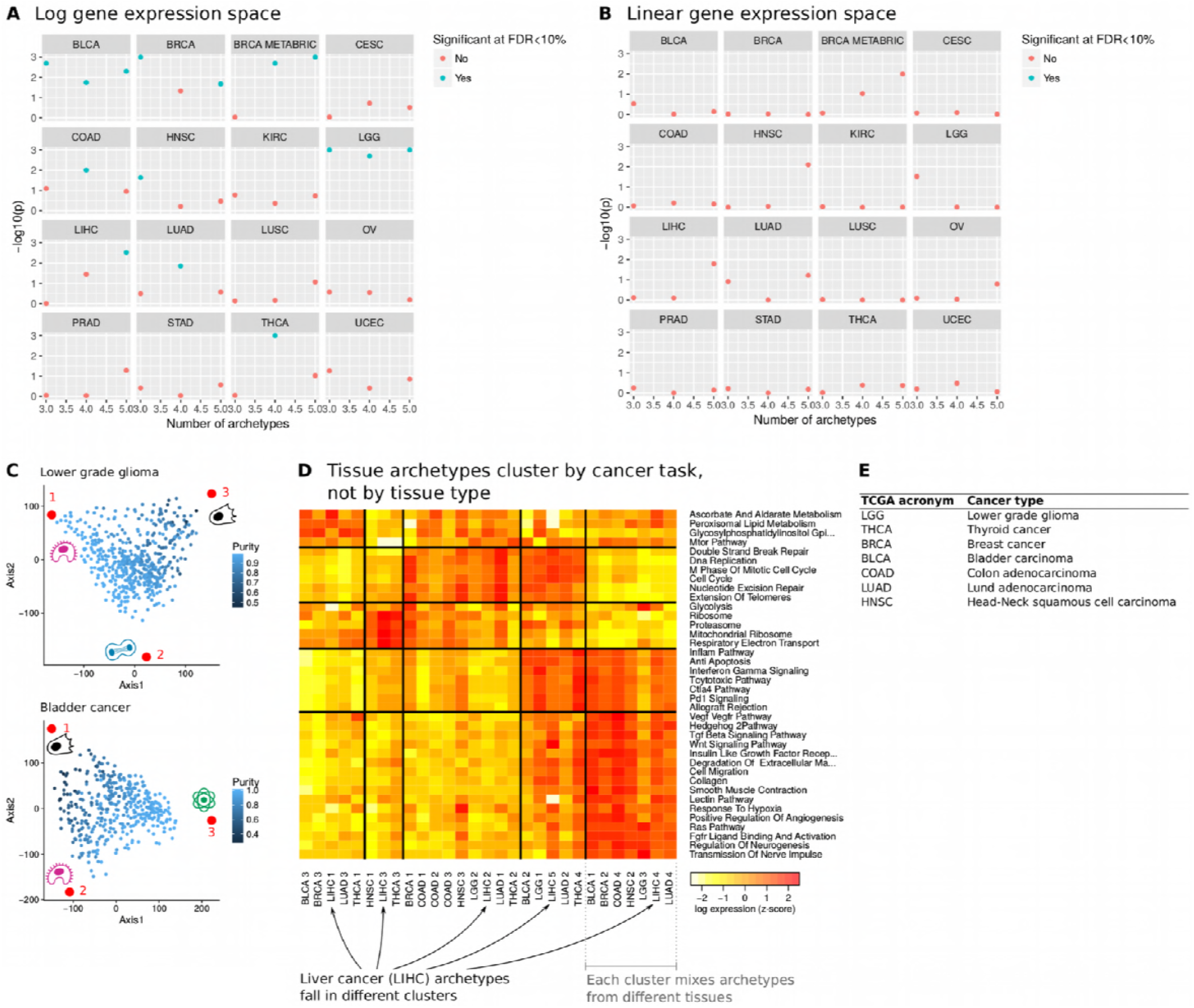
**A.** Statistical significance of polyhedra with 3, 4 and 5 archetypes in primary tumors of 15 cancer types. P-values were computed using the t-ratio test (Shoval et al. 2012; Hart et al. 2015). Polyhedra were inferred in log gene expression space. Polyhedra significant at FDR<10% appear in blue. **B.** Same as A., but for polyhedra in linear gene expression space. **C.** Purity score of tumors as computed by ESTIMATE (Yoshihara et al. 2013) as function of their position relative to the three archetypes of glioma (top) and bladder (bottom). Each blue dot represents a tumor, red dots represent archetypes. Archetype numbering corresponds to panel D and Database S1-S3. **D.** Expression of MSigDB pathways (rows) in archetypes from all cancer types (columns). Archetypes were clustered by Gaussian Mixture Model. Each cluster expresses specific MSigDB pathways. **E.** TCGA code for each cancer type.

**Fig. S2.**
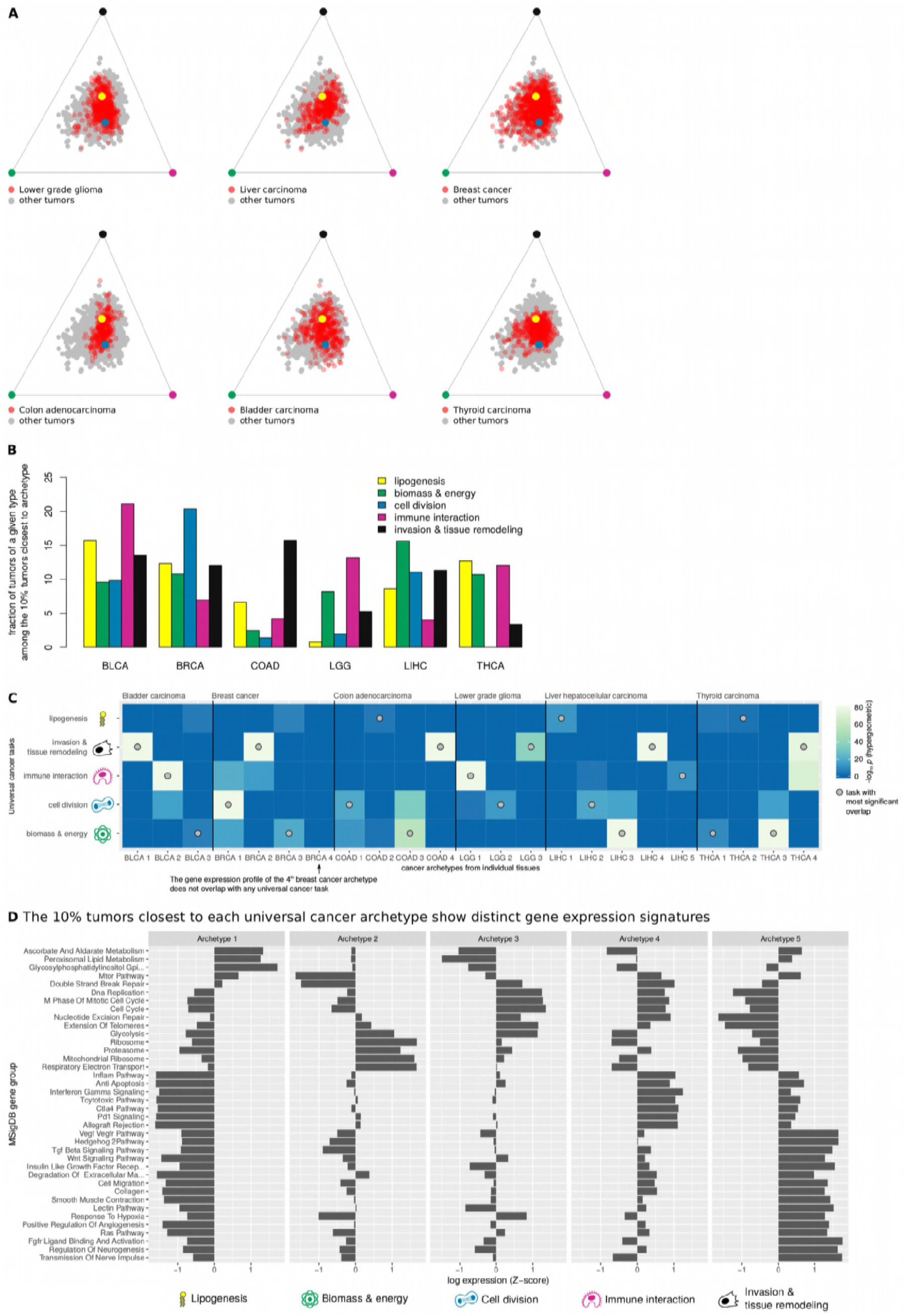
**A.** Tumors from individual tissue types spread out between several universal cancer archetypes. Red dots represent tumors from the tissue-type indicated in each panel, projected on a face of the polyhedron. The polyhedron was fitted to 3180 tumors from 6 tissue types. Grey dots represent other tumors. The remaining colored dots represent the projections of the 5 archetypes on the face. **B.** Tumors of individual tissue types are found close to multiple universal cancer archetypes. Each colored bar represents a cancer task. For example, 10.7% of thyroid tumors (THCA) are found in the 10% tumors from all cancer types closest to biomass&energy archetype. **C.** Tissue-specific archetypes can be assigned to specific universal cancer tasks with statistical significance. We compared the MSigDB pathways upregulated in each tissue-specific archetype (columns) to pathways upregulated in each universal cancer archetype (rows). The statistical similarity between pairs of archetypes was quantified using the hypergeometric test (p-values are color-coded). For each tissue-specific archetype, the most similar and statistically significant universal archetypes is signaled by a gray dot. Except in thyroid, each universal cancer task was only found once in each tissue type. For each tissue-specific archetype, there is a statistically similar universal cancer archetype, except for the 4^th^ archetype of breast cancer (HER2). **D.** Expression of MSigDB pathways in tumors close to each universal cancer archetype in gene expression space. Upregulated pathways at each archetype match tissue-specific archetype clusters (Fig. S1D) and suggest clear cancer tasks (Table 1).

**Fig. S3.**
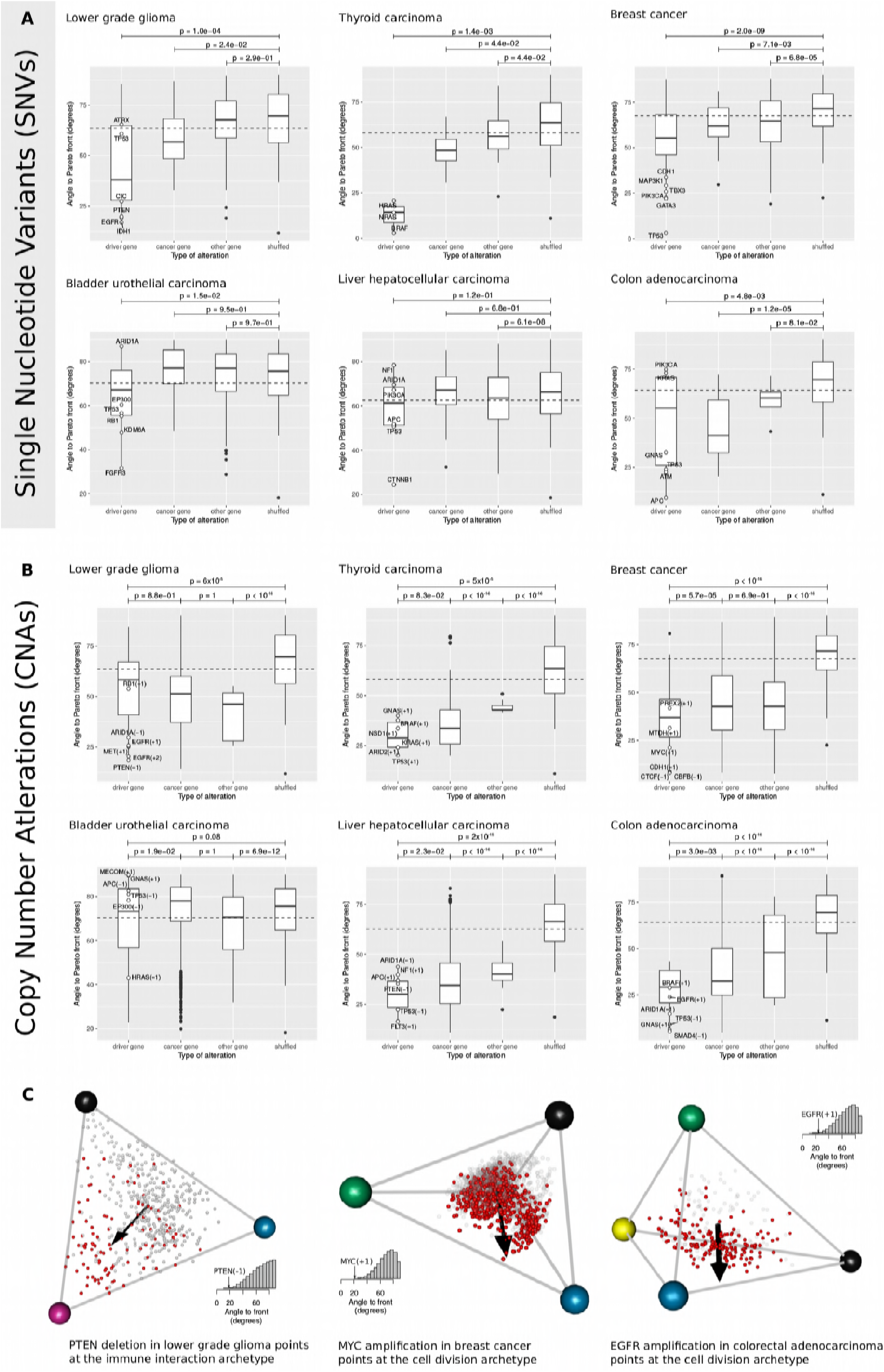
**A.** Driver SNVs are better aligned to the front of cancer than shuffled controls in glioma, thyroid, breast, bladder and colon. Shown are angle distributions of SNVs in driver genes, cancer genes (genes commonly mutated in cancer but not confidently known as drivers in this tissue), passenger genes and shuffled controls. Differences in distributions were tested using the Mann-Whitney test. **B.** Same as A, but for CNAs. **C.** In glioma, *PTEN* deletions push tumors towards the immune interaction archetype. In breast cancer, *MYC* amplification pushes tumors towards to cell division archetype. In colon cancer, *EGFR* amplification pushes tumors towards the cell division archetype.

## Additional Data table S1 (separate file)

MSigDB pathways up-regulated in the 5% tumors closest to individual archetypes. Each MSigDB pathway (feature name) is listed with the corresponding archetype. The statistical significance of the up-regulation was quantified using the Mann-Whitney U test. Uncorrected p-values appear in the table, together with median and mean difference in pathway expression of tumors in the first distance bin compared to all other tumors. MSigDB pathways only appear if statistically significant at FDR<10%, and if pathway expression is highest in tumors of the first distance bin. Results from the combined analysis of 3180 tumors from 6 cancer types appear on the sheet labeled ‘All 6 cancer types’. Also shown are individual genes up-regulated in tumors closest to individual archetypes. MSigDB pathways up-regulated when analyzing tumors from individual tissue-types separately appear in the different sheets of the spreadsheet.

## Additional Data table S2 (separate file)

Qualitative clinical features over-represented in the 5% tumors closest to individual archetypes. Each clinical feature is listed with the corresponding archetype. The statistical significance of the over-representation was quantified using the hypergeometric test. Uncorrected p-values appear in the table. Clinical features only appear if statistically significant at FDR<10%, and if the feature is most frequent among tumors of the first distance bin. Results from the combined analysis of 3810 tumors from 6 cancer types appear on the sheet labeled ‘All 6 cancer types’. Clinical features overrepresented when analyzing tumors from individual tissue-types separately appear in the different sheets of the spreadsheet. The definition of the clinical feature can be found at https://docs.gdc.cancer.gov/Data_Dictionary/viewer/.

## Additional Data table S3 (separate file)

Quantitative clinical features which take high values in the 5% tumors closest to individual archetypes. Each clinical feature is listed with the corresponding archetype. The statistical significance of the difference between tumors in the first distance bin compared to other tumors was quantified using the Mann-Whitney U test. Uncorrected p-values appear in the table. Clinical features only appear if statistically significant at FDR<10%, and if the feature has highest value in tumors of the first distance bin. Results from the combined analysis of 3180 tumors from 6 cancer types appear on the sheet labeled ‘All 6 cancer types’. Clinical features overrepresented when analyzing tumors from individual tissue-types separately appear in the different sheets of the spreadsheet. The definition of the clinical feature can be found at https://docs.gdc.cancer.gov/Data_Dictionary/viewer/.

## Additional Data table S4 (separate file)

Driver SNVs significantly aligned with the Pareto front in different cancer types (FDR<10%). For each cancer type, driver genes are listed together with the archetype / edge / face the driver points to. Also shown is the number of tumors in which the SNV was found.

## Additional Data table S5 (separate file)

Driver CNAs significantly aligned with the Pareto front in different cancer types (FDR<10%). For each cancer type, driver CNAs are listed together with the archetype / edge / face the driver points to. A CNA can be a strong deletion (−2), a weak deletion (−1), a weak amplification (+1) or a strong amplification (+2).

